# Combined knockout of *Lrrk2* and *Rab29* does not result in behavioral abnormalities *in vivo*

**DOI:** 10.1101/2020.05.13.093708

**Authors:** Melissa Conti Mazza, Victoria Nguyen, Alexandra Beilina, Jinhui Ding, Mark R. Cookson

## Abstract

Coding mutations in the *LRRK2* gene, encoding for a large protein kinase, have been shown to cause familial Parkinson’s disease (PD). The immediate biological consequence of LRRK2 mutations is to increase kinase activity, leading to the suggestion that inhibition of this enzyme might be useful therapeutically to slow disease progression. Genome-wide association studies have identified the chromosomal loci around *LRRK2* and one of its proposed substrates, *RAB29*, as contributors towards the lifetime risk of sporadic PD. Considering the evidence for interactions between LRRK2 and RAB29 on the genetic and protein levels, here we generated a double knockout mouse model and determined whether there are any consequences on brain function with aging. From a battery of motor and non-motor behavioral tests, we noted only that 18-24 month *Rab29*^-/-^ and double (*Lrrk2*^*-/-*^*/Rab29*^*-/-*^) knockout mice had diminished locomotor behavior in open field compared to wildtype mice. However, no genotype differences were seen in number of substantia nigra pars compacta (SNc) dopamine neurons or in tyrosine hydroxylase levels in the SNc and striatum, which might reflect a PD-like pathology. These results suggest that depletion of both Lrrk2 and Rab29 is tolerated, at least in mice, and support that this pathway might be able to be safely targeted for therapeutics in humans.

**Significance statement:** Genetic variation in LRRK2 that result in elevated kinase activity can cause Parkinson’s disease (PD), suggesting LRRK2 inhibition as a therapeutic strategy. RAB29, a substrate of LRRK2, has also been associated with increased PD risk. Evidence exists for an interactive relationship between LRRK2 and RAB29. Mouse models lacking either LRRK2 or RAB29 do not show brain pathologies. We hypothesized that the loss of both targets would result in additive effects across *in vivo* and post-mortem assessments in aging mice. We found that loss of both LRRK2 and RAB29 did not result in significant behavioral deficits or dopamine neuron loss. This evidence suggests that chronic inhibition of this pathway should be tolerated clinically.

## Introduction

As of 2016, 6.1 million people worldwide were diagnosed with Parkinson’s disease (PD; GBD, 2016). With an increasing aging population, the burden of PD will also rise (Dorsey & Bloem, 2018). PD is classically defined by neurodegeneration, including nigral dopamine (DA) expressing neurons, reduced motor function and non-motor symptoms, including affective and cognitive concerns. While approximately 10-15% of cases are caused by single genetic mutations, most cases have no clear cause (Genetics Home Reference, 2020) and are defined as sporadic PD. However, there is increasing evidence that genetics contributes to the etiology of sporadic PD, particularly as genome-wide association studies have identified multiple independent risk alleles that increase lifetime disease risk (Nalls et al., 2019).

Mutations in the *LRRK2* gene have been found in 1% of apparently sporadic and 4% of familial PD patients (Healy et al., 2008; Simon-Sanchez et al., 2009). Those with *LRRK2* mutations have convergent symptomatology with sporadic cases, suggesting shared pathogenic mechanisms (Healy et al., 2008; Vilas et al., 2020). LRRK2 is a large multidomain protein that contains GTP-binding Ras of complex protein (ROC), carboxyl-terminal of Roc (COR), and kinase domains (Cookson, 2010; Taylor & Alessi, 2020). Pathogenic mutations in the ROC-COR and kinase domains result in elevated kinase activity which mediates toxicity, indicating that LRRK2 inhibition may be a viable therapeutic strategy (Greggio et al., 2006; Smith et al., 2006). LRRK2 is located in the cytosol and a number of membranous structures, including the Golgi and lysosomes, and shown to affect multiple cellular functions, including autophagy, vesicular trafficking, and endolysosomal system regulation (Biskup et al., 2006; Alegre-Abarrategui et al., 2009; Beilina et al., 2014; Martin et al., 2014; Roosen & Cookson, 2016).

The *PARK16* locus is one of several chromosomal regions associated with risk of sporadic PD (Simon-Sanchez et al., 2009; Pihlstrom et al., 2015). One of the five genes in the *PARK16* locus is *RAB29* (also known as *RAB7L1*; Tucci et al., 2010; Beilina et al., 2014; Nalls et al., 2019). RAB29 belongs to the family of Rab GTPases that regulate membrane trafficking and intracellular signaling (Tucci et al., 2010; Pfeffer, 2017) and is largely localized to the Golgi complex (MacLeod et al., 2013; Beilina et al., 2014; Steger et al., 2017).

While mutations at the *LRRK2* and *RAB29* loci are independently associated with increased PD risk, there is evidence suggesting a genetic interaction between the two genes (Pihlstrom et al., 2015). Additionally, LRRK2 phosphorylates RAB29 when RAB29 is membrane and GTP bound (Steger et al., 2017; Fujimoto et al., 2018; Liu et al., 2018). PD-related LRRK2 mutations exacerbate RAB29 phosphorylation (Steger et al., 2017; Lis et al., 2018), suggesting relevance to familial PD. Additionally, RAB29 activates LRRK2 autophosphorylation and recruits LRRK2 to the Golgi and lysosomes (Beilina et al., 2014; Eguchi et al., 2018; Liu et al., 2018; Madero-Perez et al., 2018; Purlyte et al., 2018). This suggests that LRRK2 and RAB29 are reciprocal effectors in the same pathway that may play a role in membrane trafficking (MacLeod et al., 2013; Beilina et al., 2014; Eguchi et al., 2018). How the relationship between LRRK2 and RAB29 contributes to PD pathology remains unclear.

Rodent models lacking either LRRK2 or RAB29 do not exhibit any central nervous system (CNS) phenotype. No significant motor or non-motor deficits have been observed in *Lrrk2*^*-/-*^ mice or in mice with silencing of striatal LRRK2 (Volta et al., 2015). No brain pathology including DA neurodegeneration, alpha-synuclein, or ubiquitin levels were found in *Lrrk2*^*-/-*^ mice (Tong et al., 2010). Similarly, *Rab29*^*-/-*^ or *Lrrk2*^*-/-*^*/Rab29*^*-/-*^ mice did not have any DA degeneration (Kuwahara et al., 2016). These results are consistent with human genetic data where predicted loss-of-function mutations in LRRK2 are not associated with PD risk (Blauwendraat, Reed et al., 2018). Collectively, this data supports the hypothesis that LRRK2 kinase inhibition would be an effective and safe therapeutic strategy to stop human PD progression.

While these reports demonstrate a lack of CNS phenotype in single knockout models, there has, to date, been no direct behavioral comparison of *Lrrk2*^*-/-*^, *Rab29*^*-/-*^, or *Lrrk2*^*-/-*^*/Rab29*^*-/-*^ mouse models across different ages. Given the relationship between these two targets is potentially reciprocal, additive effects of double mutants might be expected. The current study aimed to fully characterize motor and non-motor assessments and DA-related pathology in *Lrrk2*^*-/-*^*/Rab29*^*-/-*^ mice compared to WT, *Lrrk2*^*-/-*^, and *Rab29*^*-/-*^ mice across different age groups. We find that loss of both Lrrk2 and Rab29 does not result in substantial behavioral deficits or loss of DA neurons, suggesting the safety of targeting this pathway therapeutically.

## Materials and Methods

### Animals

All animal procedures were performed in accordance with the Institutional Animal Care and Use Committee of the National Institute on Aging (ASP 463-LNG-2021) animal care committee’s regulations. Male and female C57BL/6J mice (N = 118; approximately 30-70 g) were used for the current study. *Rab29*^*-/-*^ mice were generated by crossbreeding *Rab29* LacZ-knockin mice (EUCOMM; Wellcome Trust Sanger Institute, Strain EM:05517) with mice expressing Cre under the constitutive CMV promoter (Jackson Laboratory, stock #006054) to remove exon 4. Single *Lrrk2*^*-/-*^ and *Rab29*^*-/-*^ mice were bred to create double knockout (*Lrrk2*^*-/-*^*/Rab29*^*-/-*^) mice. Genotypes were confirmed using gel-based PCR. The mice were provided with group housing and given access to food and water ad libitum and housed in a facility with 12hr light/dark cycles with temperature maintained at 22-23°C, and experiments took place between Zeitgeber Time ZT2 – ZT11.

### Experimental design

Mice were tested in three age groups to determine genotype- and age-related differences across multiple behaviors (see Figure 1 for *n* by age and genotype). Mice were handled prior to behavioral testing to reduce stress response and body weight was recorded. For 2 weeks, mice were tested daily on various behavioral tasks using the Rodent Behavior Core facility (National Institute of Mental Health). These tasks included: open field, elevated plus maze, spontaneous alternation, buried treat test, grip strength, and accelerating rotarod. Experimenters evaluating behavior were blinded to age and genotype. Once behavioral testing was complete, at 18-24 months of age, mice were weighed and tissue was collected for postmortem analyses. The anterior brain was flash frozen and stored at -80°C for striatal tissue dissection for western blot analysis. The posterior brain was post-fixed in 4% paraformaldehyde for 48hr and then stored in 30% sucrose prior to tissue preparation for immunohistochemistry.

**Figure 1.**
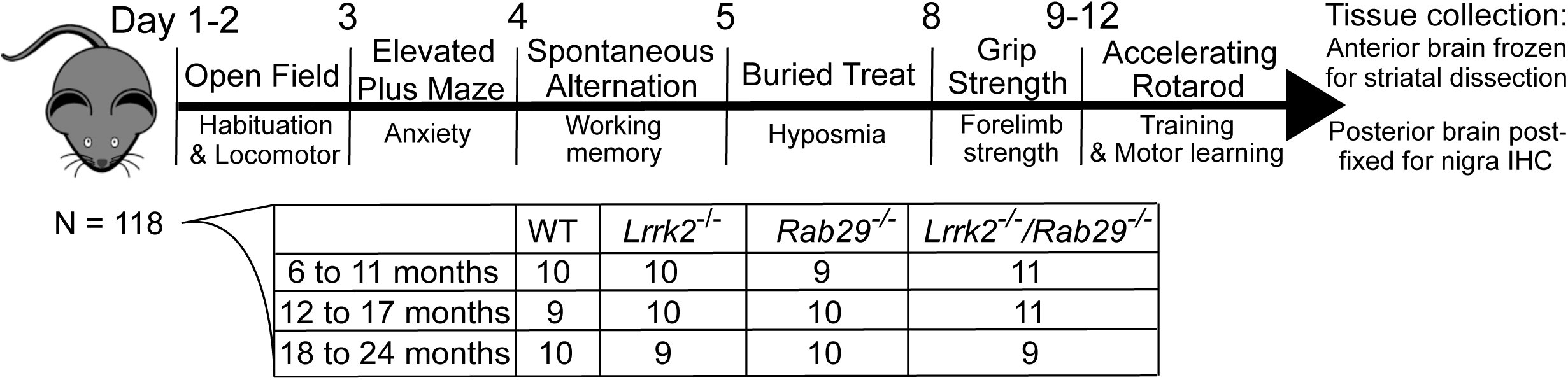
Study design. Male and female mice were handled prior to behavioral testing. Mice underwent all testing from 8:00 AM - 5:00 PM during their light cycle. Once all behavioral testing was complete, brain tissue was collected for post-mortem analysis. The anterior brain was flash frozen on dry ice for striatal tissue dissection and western blot analysis. The posterior brain was post-fixed in 4% paraformaldehyde for 48 hours then stored in 30% sucrose for immunohistochemistry staining of the SNc.

### Behavioral tests

#### Open field

Mice underwent open field testing twice; first to habituate to the test and the second to test for baseline locomotor performance. Each day, mice were placed in the testing room under red light for 1hr prior to habituate to the room. Mice were then placed in a plexiglass chamber equipped with 2 layers of beams (4 × 8; Photobeam Activity System-Open Field, San Diego Instruments) for a 30 min trial and allowed to explore freely. Ambulatory frequency, a measure for repetitive, stereotypic movements, was calculated as the ratio of ambulatory movements to the sum of unique beam breaks by ambulatory movement. Path length, a measure for total distance travelled, was calculated through relative distance by coordinates of beam breaks. All other data are measured as counts. Only data from the test day is shown.

### Elevated plus maze

The elevated plus maze was used to measure differences in anxiety-related behaviors (Komada, Takao, & Miyakawa, 2008). Mice were placed in testing room lit at 100 lux and allowed to habituate for 1hr. Mice were then placed in center of elevated plus maze containing 2 open and 2 closed arms 77.5 cm from the floor (each arm measured 38.1 × 5.1 cm with walls 15.2 cm tall). Mice started off facing a closed arm and allowed to explore the maze freely for 10 min. Trials were recorded and later scored using Top Scan 3.0 software. Anxiety-related measures were determined as percent open arm behaviors were the sum of the open arms was divided by the sum of all the arms.

### Spontaneous alternation

To detect deficits in working memory, spontaneous alternation was carried out on a plexiglass Y-maze with each arm measuring 40 × 8 x 12 cm (Kimura, Devi, & Ohno, 2010). Each mouse was placed in the center of the maze and allowed to explore the arms freely for 8 min. The sequence of arm entry and total number of arm entries was recorded. Spontaneous alternation (percent) was determined by taking the total number of consecutive arm entries divided by the number of possible alternations (total arm entries minus 2) multiplied by 100. Mice with less than 10 arm entries were excluded from this analysis (*n*=13).

### Buried treat test

To determine whether mice exhibited hyposmia, a promising marker for PD severity (Roos et al., 2019), mice were tested on the buried treat test (Dranka et al., 2014). Three days prior to testing, mice were habituated to fruit-scented cereal treats. The day before testing, mice were food deprived for a minimum of 12hr. On test day, mice were individually placed in a new cage with 3cm of clean bedding. A chow pellet and cereal treat were buried 0.5cm deep. The mouse was free to locate the treat and time to treat was recorded with a maximum time of 5min. Three trials were performed with bedding replaced in between each trial. The average time across trials was recorded for each mouse.

### Grip strength

The grip strength test was used to quantify differences in skeletal muscular strength in mouse forelimbs (Luszczki & Czuczwar, 2007). The grip strength apparatus (BioSeb) includes a wire grid (8 × 8 cm) where mice were lifted by the tail so that the forepaws grasped the grid and then were gently pulled backward by the tail until they released the grid. The maximal force was recorded and the mean of 5 trials was included for each mouse.

### Accelerating rotarod

Motor performance and motor learning were testing using the accelerating rotarod assay. Mice were habituated to the test by walking on the rod for 1 min at the stable speed of 5 rpm. Mice were then tested for the following 3 days where they underwent 3 trials each day at an increasing speed from 4 – 40 rpm across 5 min. Trials were separated by at least 15 min. Latency to fall was recorded for each trial and the mean across trials was entered for each day.

### Postmortem analyses

#### Immunohistochemistry

Post-fixed posterior brains were sectioned on a microtome into 30 µm coronal sections and stored in cryoprotectant at -20°C as previously described (Hauser et al., 2017). Briefly, sections containing the substantia nigra pars compacta (SNc) were quenched in glycine (0.3 M in PBS) and were then washed in PBS 3 times. Sections were blocked for 1 hr at RT in blocking buffer (1% BSA, 0.3% Triton X-100, and 10% donkey serum in PBS). Following 3 PBS rinses, sections were incubated in primary antibodies at 4°C overnight: tyrosine hydroxylase (TH, Abcam ab112, rabbit polyclonal, 1:500) and NeuN (Abcam, ab104224, mouse monoclonal, 1:1000) in 1% BSA, 0.3% Triton X-100, and 1% donkey serum in PBS. Sections were rinsed and incubated in secondary antibodies for 1 hr at RT (Alexa Fluor 488 donkey anti-rabbit IgG, 1:500; Alexa Fluor 568 donkey anti-mouse IgG, 1:500; Hoechst 33,342, 1:10,000). Sections were rinsed and mounted on glass slides which were then rinsed in water and coverslipped using Prolong Gold mounting media. Slides were imaged on a Zeiss 880 confocal microscope at 10x magnification. Fluorescence intensity was quantified for each nigra section using ImageJ. Data were recorded as the average of values across 3 nigra sections for a mean fluorescence intensity per mouse.

### Western blot

Western blot procedures were carried out as previously described (Roosen et al., 2020). Frozen striatal tissue was dissected and homogenized in lysis buffer (20 mM Tris pH 7.5, 10% glycerol, 1 mM EDTA, 150 mM NaCl, 1x protease inhibitor cocktail (Halt), 1x phosphatase inhibitor cocktail (Halt)) and lysed on ice for 30 min. Samples were spun at 20,000 g for 10 min at 4°C and protein supernatant was separated. Protein concentrations were determined by the Pierce 660 assay. Protein lysates were diluted to 10 µg per sample and boiled in 1x Laemli Sample Buffer (Bio-Rad). Samples were loaded on pre-cast 4-20% TGX polyacrylamide gels (Criterion, Bio-Rad). Electrophoresis was performed in 1x pre-mixed buffer (10 mM Tris, 10 mM Tricine, 0.01% SDS, pH 8.3, diluted with water) using the Criterion Vertical Electrophoresis Cell (Bio-Rad). Gels were then transferred to 0.45 µm pore-size nitrocellulose membranes (Bio-Rad) using the Trans-Blot Turbo Transfer System (Bio-Rad). Membranes were blocked using a 1:1 solution of phosphate buffered saline (PBS) and Odyssey Blocking Buffer (LiCor). After blocking, membranes were incubated overnight on gentle agitation at 4°C with primary antibodies (TH, 1:1000; GAPDH, Sigma, G8795, mouse monoclonal, 1:10,000) diluted in antibody buffer (1:1 of Tris buffered saline (TBS) with 0.1% Tween and Odyssey Blocking Buffer. Membranes were then washed 3 times for 5 min each in TBS-0.1% Tween followed by a 1 hr incubation at room temperature in fluorescent secondary antibodies (IRDye, Li-Cor) diluted 1:15,000 in antibody buffer. After 3 additional washes, western blots were imaged using the Odyssey CLx system (Li-Cor) and quantified using Image Studio software.

### Statistical analyses

Divergence from a normal distribution was estimated using the Shapiro-Wilk Normality test. Accelerating rotarod data was analyzed using a three-way mixed design ANOVA (genotype x age x test day). Postmortem analyses were analyzed for genotype differences in the 18 – 24 month age group using a one-way between-subjects ANOVA. All other data was analyzed with a two-way between-subjects ANOVA (genotype x age). Tukey *post-hoc* analyses were used to determine significant differences when appropriate. All analyses were carried out using RStudio (https://rstudio.com/) and alpha was set to 0.05. For one-way ANOVAs and main effects, Cohen’s f and achieved power were determined using the “powerAnalysis” package in R. For interactions, eta squared was calculated using the “rstatix” package in R. Eta squared was converted to Cohen’s f to determine achieved power using GPower (3.0.10; <u>https://www.psychologie.hhu.de/arbeitsgruppen/allgemeine-psychologie-und-arbeitspsychologie/gpower.html</u>). Calculated effect sizes (Cohen’s f), *n*, and *p* values calculated from ANOVAs were used when determining achieved power (see Table 1 for values).

**Table 1.** Statistical table. *Post-hoc* power calculations for each statistical test were recorded. To calculate achieved power, we used the calculated effect size, group sample size, and the *p* value from the indicated test. Superscript letters in the text refer to the first column of the table.

|  | Data Structure | Type of Test | Effect Size | Power |
| --- | --- | --- | --- | --- |
| a | Non-normal distribution | Two-way ANOVA | 0.351 | 0.790 |
| b | Non-normal distribution | Two-way ANOVA | 0.357 | 0.786 |
| c | Normal distribution | One-way ANOVA | 0.746 | 0.919 |
| d | Normal distribution | One-way ANOVA | 0.905 | 0.969 |
| e | Non-normal distribution | One-way ANOVA | 0.702 | 0.901 |
| f | Non-normal distribution | Three-way ANOVA | 1.426 | 0.999 |
| g | Non-normal distribution | Three-way ANOVA | 0.417 | 0.999 |
| h | Normal distribution | Two-way ANOVA | 0.327 | 0.999 |
| i | Non-normal distribution | Two-way ANOVA | 0.314 | 0.999 |
| j | Non-normal distribution | Two-way ANOVA | 0.467 | 0.999 |
| k | Normal distribution | Two-way ANOVA | 0.415 | 0.720 |
| l | Non-normal distribution | Two-way ANOVA | 0.328 | 0.715 |
| m | Non-normal distribution | Two-way ANOVA | 0.366 | 0.795 |

## Results

### Motor behaviors

Age- and gene-related differences in baseline locomotor behavior was determined by placing WT, *Lrrk2*^*-/-*^, *Rab29*^*-/-*^, and *Lrrk2*^*-/-*^*/Rab29*^*-/-*^ knockout mice in open field chambers. There were no differences due to age or genotype in total activity, fine movement, rearing activity, ambulatory activity, ambulatory frequency, and path length (Fig. 2A – F). A significant genotype x age interaction was noted for ambulatory count (F_6,106_ = 2.23, p=0.045^a^) and path length (F_6,106_ = 2.32, p=0.038^b^); however, no *post-hoc* comparisons between groups were significant. In the oldest age group (18 – 24 months), there was a main effect of genotype where *Rab29*^*-/-*^ and *Lrrk2*^*-/-*^*/Rab29*^*-/-*^ mice had lower ambulatory count (one-way ANOVA, F_3,34_ = 4.05, p=0.015^c^; Fig. 2D, p<0.05) and travelled less according to path length (F_3,34_ = 5.10, p=0.005^d^; Fig. 2F, p<0.05). Furthermore *Lrrk2*^*-/-*^*/Rab29*^*-/-*^ knockout mice had reduced ambulatory frequency compared to WT (F_3,34_ = 3.74, p=0.02^e^; Fig. 2E, p<0.05).

**Figure 2.**
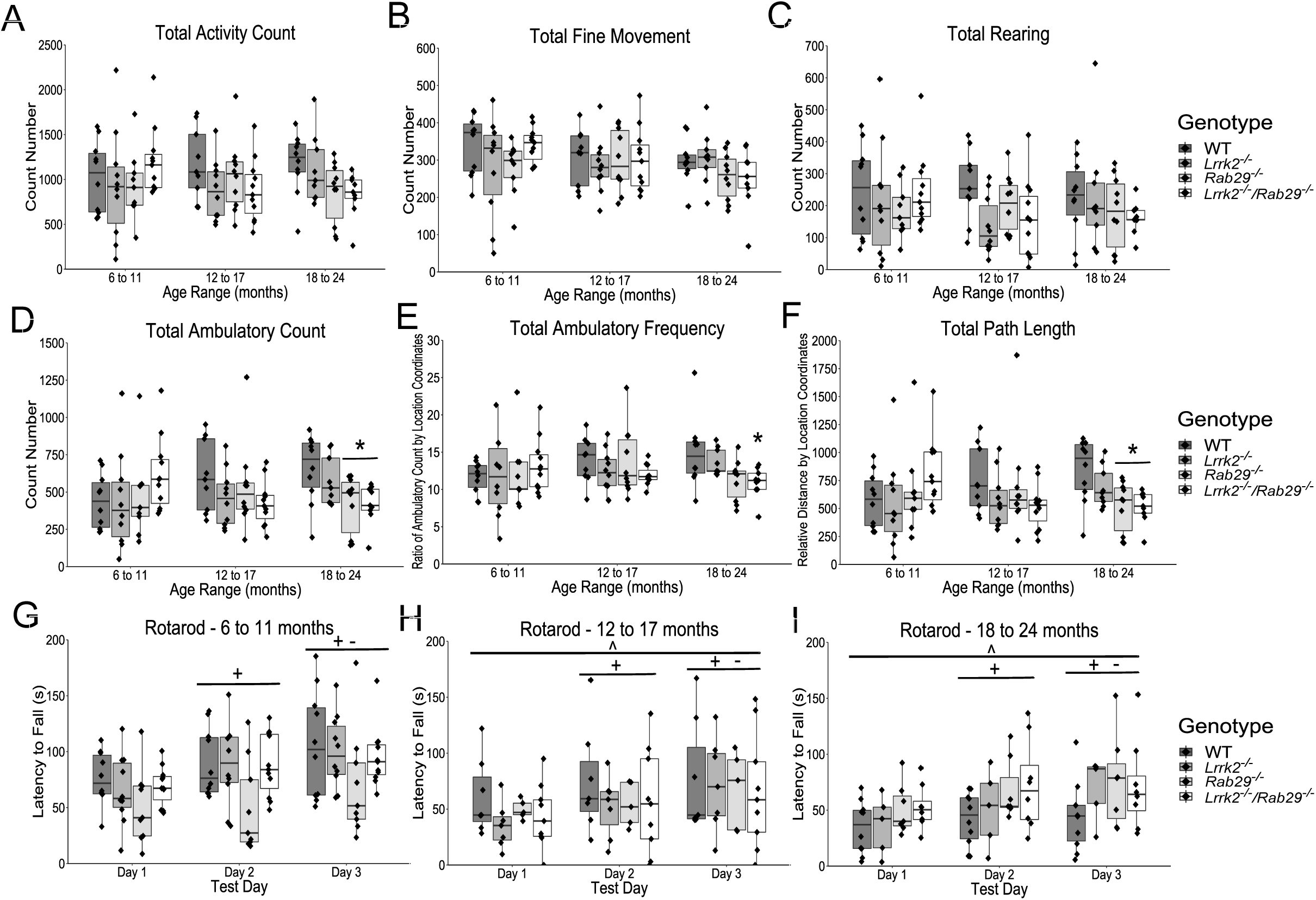
Locomotor behavior characterization of aging single and double knockout mice. Open field measurements of locomotor activity across total session including; activity count (A), fine movement count (B); rearing count (C); ambulatory activity count (D); ambulatory frequency (E; stereotypy measure); and path length (F). Accelerating rotarod to show differences in motor performance and learning for each age: 6 to 11 (G), 12 to 17 (H), and 18 to 24 month (I). * p<0.05 vs. WT 18 to 24 month, ^ p<0.05 vs. 6 to 11month, + p<0.05 vs. Day 1, and - p<0.05 vs. Day 2.

Motor performance was also measured using accelerating rotarod. There was a main effect of day (F_2,172_ = 57.66, p=2×10^−16f^), where latency to fall significantly increased each day (p<0.05). There was also a main effect of age where the youngest group (6 – 11 months) had greater latencies than the two older age groups (F_2,172_ = 6.36, p=0.003^g^; p<0.05).

Overall, the motor evaluations indicated a decline in some open field measures aged *Lrrk2*^*-/-*^*/Rab29*^*-/-*^ mice but no effects of genotype on motor learning as determined by rotarod. These results indicate that motor function is impacted by combined Lrrk2 and Rab29 deficiency in modest ways.

### Non-motor behaviors

To further characterize the effects of Lrrk2 and Rab29 deficiency, mice were tested on a variety of non-motor behaviors. The elevated plus maze was used to measure anxiety-related behaviors. A main effect of age was found where 6 – 11 month mice had the lowest amount of anxiety as measured by more open arm entries (F_2,105_ = 5.06, p=0.008^h^; Fig. 3A, p=0.006) and longer time spent in the open arms (F_2,105_ = 4.73, p=0.012^i^; Fig. 3B, p=0.008) compared to 18 – 24 month mice as well as longer distance travelled in open arms compared to 12 – 17 and 18 – 24 month mice (F_2,105_ = 9.41, p=0.0002^j^; Fig. 3C, p<0.05). A significant genotype x age interaction was identified for percent open arm entries (F_6,105_ = 3.33, p=0.005^k^) and distance travelled (F_6,105_ = 2.26, p=0.043^l^). *Rab29*^*-/-*^ mice aged 6 – 11 months entered open arms more than 6 – 11 month *Lrrk2*^*-/-*^*/Rab29*^*-/-*^ mice and 18 – 24 month *Rab29*^*-/-*^ mice (Fig. 3A, p<0.05). *Lrrk2*^*-/-*^ mice aged 6 – 11 months travelled farther in open arms than 12 – 17 and 18 – 24 month *Lrrk2*^*-/-*^ mice and 6 – 11 month *Lrrk2*^*-/-*^*/Rab29*^*-/-*^ mice (Fig. 3B; p<0.05).

**Figure 3.**
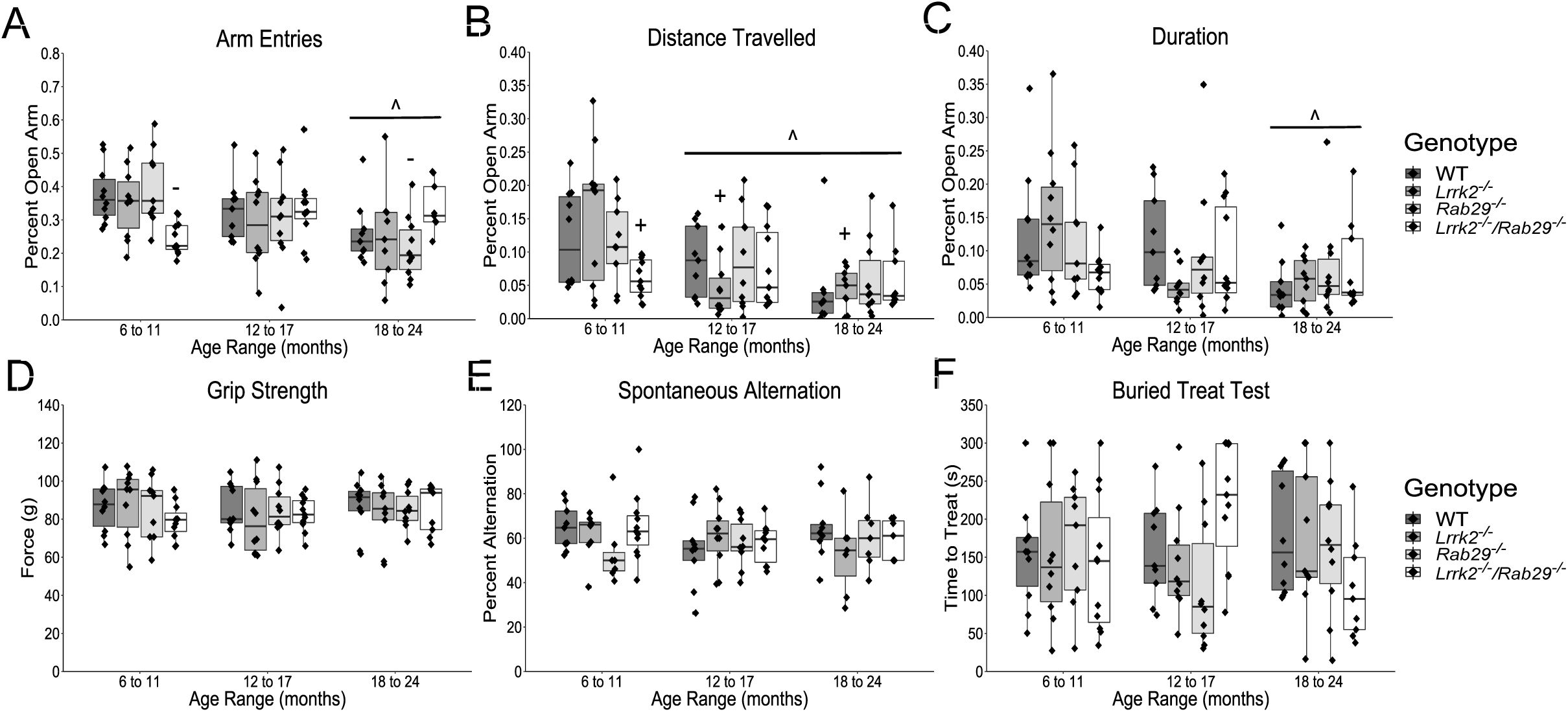
Nonmotor behavior characterization of aging single and double knockout mice. Elevated plus field measurements of anxiety determined by percent of open arm; entries (A), distance travelled (B); and duration (C). Other nonmotor assays included; grip strength (D); spontaneous alternation for working memory (E); and buried treat test as a measure for hypsomia (F). ^ p<0.05 vs. 6 to 11 month, + p<0.05 vs. Lrrk2: 6 to 11 month, and - p<0.05 vs. Rab29: 6 to 11 month.

There were no reported age- or genotype-related differences on forelimb strength as measured by the grip strength test (Fig. 3D) or working memory as measured by spontaneous alternation (Fig. 3E). There was a genotype x age interaction found for the buried treat test measurement of hyposmia (F_6,106_ = 2.39, p=0.034^m^); however, no significant *post-hoc* comparisons were noted between genotypes (Fig. 3F). Overall, the only non-motor behaviors to show group differences were anxiety-related and while 6 – 11 month *Lrrk2*^*-/-*^*/Rab29*^*-/-*^ mice exhibited greater anxiety behavior compared to single knockout mice, age was the most significant factor in increased anxiety behavior.

### Postmortem analyses

Due to the reduced motor activity observed in 18 – 24 month *Rab29*^*-/-*^ and *Lrrk2*^*-/-*^*/Rab29*^*-/-*^ mice, postmortem analyses focused on genotypic differences in the oldest age group. Using immunohistochemistry, the SNc was stained for tyrosine hydroxylase (TH; the rate limiting enzyme in dopamine synthesis) and neuronal nuclei (NeuN) to determine any loss of dopamine cell bodies. There were no differences among genotypes in nigral TH or NeuN fluorescence intensity (Fig. 4A-C).

**Figure 4.**
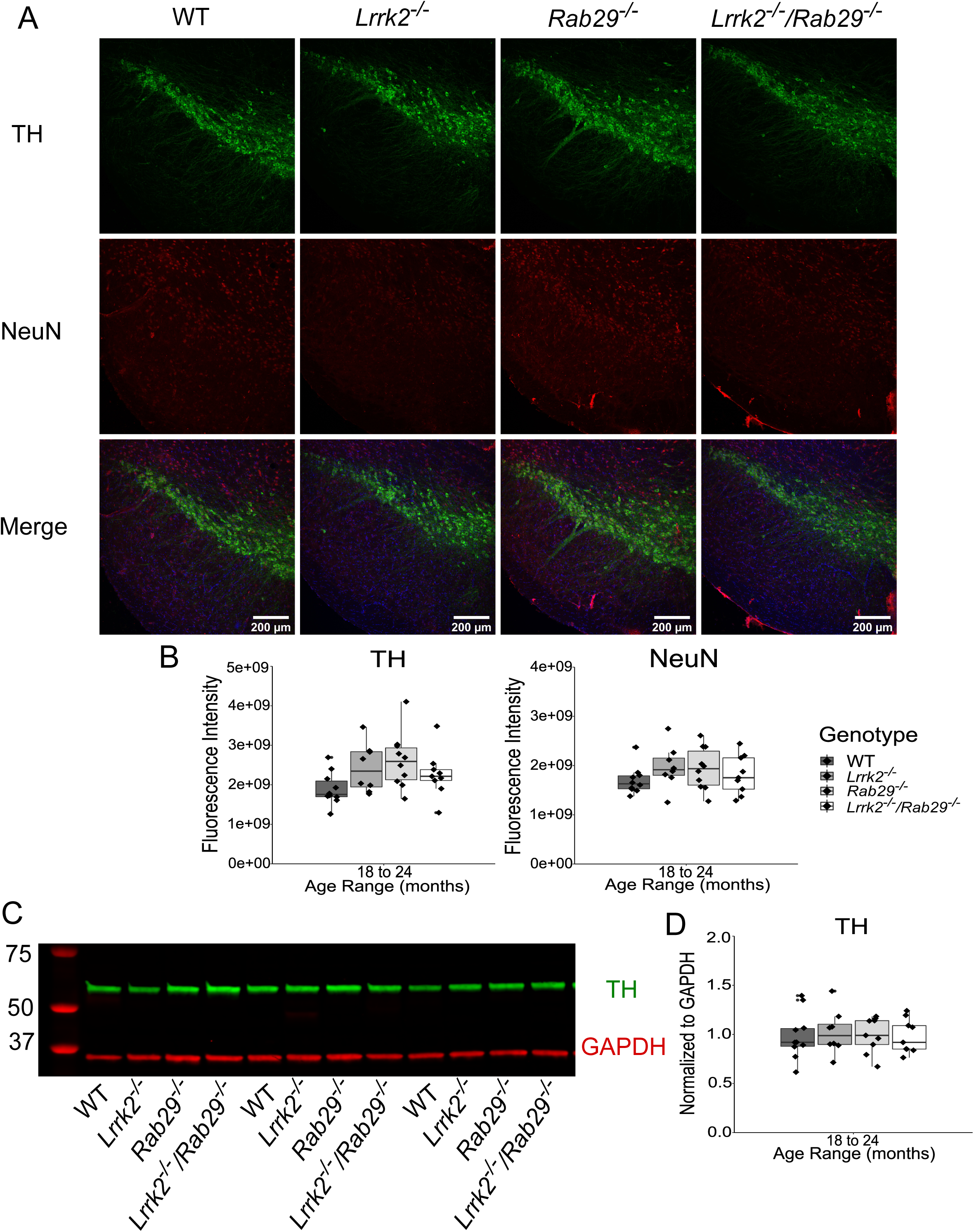
Postmortem analyses of dopamine loss in the SNc and striatum of 18 – 24 month single and double knockout mice. Immunohistochemistry images of SNc stained for TH (green), NeuN (red), and Hoechst (blue; A). Genotype differences in TH (B) and NeuN (C) mean fluorescence intensity were quantified. Western blot analysis of striatal tissue expression of TH and GAPDH (D). Striatal TH was normalized to GAPDH and quantified for genotype differences (E).

Loss of TH protein in the striatum could indicate degeneration at the level of dopamine projections from the SNc. Striatal tissue was isolated from fresh frozen anterior brain and analyzed for TH expression by western blot. There were no differences among genotypes in striatal TH, normalized to GAPDH as loading control, expression (Fig. 4D-E). Overall, this suggests that there was no pathological neurodegeneration observed in dopamine neurons at the level of the cell body in the SNc or striatal projections.

## Discussion

Prior literature suggests that *LRRK2* and *RAB29* operate within the same genetic pathway (Purlyte et al., 2018; Gomez et al., 2019; for review, see Taylor & Alessi, 2020). LRRK2 can phosphorylate membrane bound RAB29, which is normally resident at the trans-Golgi network (TGN; Steger et al., 2017; Fujimoto et al., 2018; Gomez et al., 2019). This interaction indicates that RAB29 is a downstream effector of LRRK2, where it may mediate alterations in membrane trafficking and neurite outgrowth (Kuwahara et al., 2016; Feng et al., 2018). Consistent with these ideas, hyperactive mutations in LRRK2 enhance RAB29-dependent changes in TGN morphology (Beilina et al., 2014). However, overexpression of RAB29 can activate LRRK2, suggesting an upstream regulatory relationship of the GTPase on kinase activity (Beilina et al., 2014; Liu et al., 2018; Purlyte et al., 2018). In addition, RAB29 and LRRK2 can affect lysosomal morphology in HEK cells (Eguchi et al., 2018). Therefore, although it is clear that RAB29 and LRRK2 have related functions in cells, there are several uncertainties for direction of effect and the downstream outputs of this pathway (for review, see Kuwahara & Iwatsubo, 2020).

To investigate the nature of the interaction between these PD-related targets *in vivo*, we generated a LRRK2 and RAB29 double knockout (*Lrrk2*^*-/-*^*/Rab29*^*-/-*^) mouse model. A baseline characterization of PD-related behavior and pathology was performed to determine whether loss of both LRRK2 and RAB29 would result in brain phenotypes. No differences between genotypes were seen in measures of strength, working memory, or hyposmia. We noted a mild motor phenotype in the *Rab29*^*-/-*^ and *Lrrk2*^*-/-*^*/Rab29*^*-/-*^ animals, which we interpret to be dependent on RAB29, not LRRK2. Additionally, when looking specifically at the oldest age group (18-24 month), no differences in genotype were observed in DA-related pathology in the SNc and striatum. The lack of robust behavioral phenotype and DA degeneration is consistent with previous reports on single LRRK2 and Rab29 knockout models (Tong et al., 2010; Volta et al., 2015; Kuwahara et al., 2016). The lack of CNS phenotypes unfortunately prevents an evaluation of the pathway relationship between LRRK2 and RAB29 at this time.

Elevated kinase activity associated with pathogenic LRRK2 mutations (Greggio et al., 2006; Smith et al., 2006) as well as increased LRRK2 phosphorylation associated with sporadic PD indicate that LRRK2 inhibition is a potential therapeutic strategy for PD patients (Di Maio et al., 2018). Given the long duration and gradual nature of PD, chronic, ongoing treatment will be required and it is therefore important to understand the consequences of dampening LRRK2 kinase activity on a long-term basis. Information from knockout models and loss-of-function mutant carriers may grant some clarity towards this issue. As shown here, knocking out Lrrk2 and its interactor Rab29 had no significant effect on DA pathology related to PD in the aging mouse model, consistent with prior reports of single LRRK2 knockouts (Volta et al., 2015; Kuwahara et al., 2016). However, double knockout of LRRK2 and LRRK1 is reported to cause early mortality and significant DA degeneration highlighting the importance of selectively targeting LRRK2 (Giaime et al., 2017).

Overall, our results support the concept that the LRRK2-RAB29 pathway can be safely targeted for PD-related therapeutics, but suggest that LRRK2 remains the better target compared to RAB29 due to the mild motor phenotypes seen in the *Rab29*^*-/-*^ and *Lrrk2*^*-/-*^*/Rab29*^*-/-*^ animals. However, the lack of a robust brain phenotype precludes us from answering the question of the directional relationship between LRRK2 and RAB29 with the data presented here. There are peripheral phenotypes in models of LRRK2 deficiency, such as accumulation of lysosomal markers in the kidney and invasion of the alveolar space by type-II pneumocytes in the lung (Baptista et al., 2013; Kuwahara et al., 2016; Whiffin et al., 2019). Further studies on the peripheral phenotypes in the *Rab29*^*-/-*^ and *Lrrk2*^*-/-*^*/Rab29*^*-/-*^ animals will be required to clarify whether LRRK2 is upstream or downstream of RAB29 *in vivo*.

